# Unveiling Driver Modules in Lung Cancer: A Clustering-Based Gene-Gene Interaction Network Analysis

**DOI:** 10.1101/2023.11.01.565104

**Authors:** Golnaz Taheri, Marcell Szalai, Mahnaz Habibi, Panagiotis Papapetrou

## Abstract

Lung cancer, which is the leading cause of cancer-related death worldwide and is characterized by genetic changes and heterogeneity, presents a significant treatment challenge. Existing approaches utilizing Machine Learning (ML) techniques for identifying driver modules lack specificity, particularly for lung cancer. This study addresses this limitation by proposing a novel method that combines gene-gene interaction network construction with ML-based clustering to identify lung cancer-specific driver modules. The methodology involves mapping biological processes to genes and constructing a weighted gene-gene interaction network to identify correlations within gene clusters. A clustering algorithm is then applied to identify potential cancer-driver modules, focusing on biologically relevant modules that contribute to lung cancer development. The results highlight the effectiveness and robustness of the clustering approach, identifying 110 unique clusters ranging in size from 4 to 10. These clusters surpass evaluation requirements and demonstrate significant relevance to critical cancer-related pathways. The identified driver modules hold promise for influencing future approaches to lung cancer diagnosis, prognosis, and treatment. This research expands our understanding of lung cancer and sets the stage for further investigations and potential clinical advancements.

## 1 Introduction

Lung cancer stands as the foremost contributor to cancer-related mortality globally, accounting for an estimated 1.8 million deaths in 2020, which represents almost 18.4% of all cancer deaths [1]. Unfortunately, the prognosis for lung cancer remains unfavorable, with only 18.6% of patients surviving beyond five years [2]. Smoking is responsible for the majority of cases, while a notable percentage occurs in non-smokers [3, 4]. Environmental factors like asbestos, radon gas, and pollution also contribute to lung cancer [5]. There are two main types: small cell lung cancer (SCLC) and non-small cell lung cancer (NSCLC), with NSCLC being the most common and having high incidence and mortality rates [6]. In 2020 an estimate of 2,206,771 new cases of NSCLC was reported, while the mortality rate for this cancer in the same year reached 1,796,144 individuals. NSCLC is further categorized into adenocarcinoma, squamous cell carcinoma, and large cell carcinoma [7]. Genomic studies have revealed the complex molecular landscape associated with lung tumors influenced by factors, such as smoking and environmental exposures [7]. Advancements in deep sequencing technologies, exemplified by projects like The Cancer Genome Atlas (TCGA) [9], have generated extensive cancer genomics data. This wealth of information presents unprecedented opportunities to comprehend the mechanisms of carcinogenesis. Identifying cancer-related somatic mutations, driver modules, and functionally interconnected gene alterations is crucial for understanding cancer initiation, progression, and development. Driver modules actively contribute to cancer’s hallmark characteristics and uncovering them enhances our understanding of cancer and aids in designing effective treatment strategies [10].

Various computational approaches have emerged to prioritize genes independently and identify candidate driver genes, which are genes causally linked to different diseases and oncogenesis [11–14]. These methods combine somatic mutation data with additional information, such as interaction networks or gene expression data, to generate gene rankings. While these rankings offer valuable insights into potential genes of interest, it is important to note that mutations at different genomic loci can contribute to the same disease [15]. This genetic variation might point to an underlying biological process in which genes linked to cancer could operate as possible driver modules and functional pathways. For the purpose of finding such candidate modules, several computational techniques have been developed. [16, 17].

### Related Work

Mutual exclusivity refers to the non-random occurrence of genomic alterations without co-occurrence. MEMo is a computational method introduced by Ciriello et al. [18] to identify mutually exclusive alteration sets. MEMo compares observed co-occurrence frequencies with a null model, considering alteration frequencies and sample cohort size. It successfully highlights mutually exclusive modules and identifies promising genes selectively altered in tumors. However, MEMo relies on existing biological knowledge and stringent filters, limiting its ability to discover associations between less-studied genes. While useful for evaluating mutation combinations, mutual exclusivity has limitations. It does not encompass all possibilities, as concurrent driver mutations can occur and the pattern applies only within the same pathway, not across different pathways. Therefore, mutual exclusivity alone is inadequate for characterizing all functional mutation combinations. Detecting cancer-driving mutations among infrequently mutated genes is challenging. Cho et al. [19] introduced MUFFINN, a pathway-focused technique that considers mutations in both individual genes and their functional network neighbors. MUFFINN demonstrates increased sensitivity compared to gene-focused analyses, making it valuable for prioritizing cancer genes. Importantly, it remains effective even with smaller patient populations, enhancing cancer genome projects. Advanced computational techniques utilize biological pathways and networks to identify cancer driver mutations and pathways. De novo methods analyze somatic mutation patterns across tumors to infer cancer-related genes and pathways. Dimitrakopoulos et al. [20] discuss recent advancements, limitations, and future challenges in detecting cancer genes and pathways, emphasizing the need for de novo approaches to overcome biases in prior knowledge-based methods. Pathway-based methods rely on reference databases, while network-based methods favor well-studied genes. In contrast, techniques identifying combinatorial mutation patterns learn pathways from scratch, enhancing sensitivity by managing noise and evaluating statistical uncertainties. Zhang et al. [21] developed an integrated approach that combines mutual exclusivity, coverage, and protein-protein interaction data to construct an edge-weighted network. It outperformed other methods in identifying known cancer driver modules and demonstrated high precision in distinguishing normal and tumor samples using pan-cancer data.

These studies have a limitation in their broad-spectrum approach to cancer, which lacks specificity for particular types of cancer like lung cancer. While the findings are generally promising, their applicability to specific cancers, including lung cancer, remains untested and uncertain. The success observed in these studies may not necessarily translate to equally successful outcomes when applied to lung cancer or other specific types of cancer. This limitation highlights the importance of further research that focuses on individual cancer types to gain a more nuanced understanding of the disease.

### Contributions

This research introduces a novel approach for identifying statistically significant gene clusters using a graph-based representation of gene interactions. It addresses limitations in current broad-spectrum cancer studies and focuses specifically on lung cancer. By utilizing clustering techniques, this study aims to identify unique driver modules specific to lung cancer, providing valuable insights at the molecular level. This targeted method enhances personalized treatment strategies and bridges the gap between general and specific cancer research, improving the precision and relevance of findings for lung cancer patients. The main contributions of this work are summarized as follows:

− **Novelty**. This study introduces a novel approach to overcome limitations in current broad-spectrum cancer studies. Existing research lacks specificity, making it uncertain for specific cancer types like lung cancer. This study focuses on identifying unique driver modules in lung cancer using clustering, providing a more targeted understanding at the molecular level and may lead to more reliable insights and personalized treatment strategies.
− **Significance**. The identification of statistically significant gene clusters provides a fresh perspective on gene-based indicators of cancer, indicating that modules of various sizes can contribute to lung cancer development. Additionally, identifying commonly occurring genes across multiple clusters helps identify significant contributors to lung cancer.
− **Clinical investigation**. The study’s findings have practical implications for understanding the genetic landscape of lung cancer. This knowledge can impact diagnostics, prognosis, and the development of targeted therapies, enhancing our understanding of the role of genes and modules in lung cancer and their implications for therapeutic interventions.

## 2 A Workflow for Clustering-Based Gene-Gene Interaction Network Analysis

In this section, we introduce a workflow for identifying driver modules in lung cancer, which involves two key steps. Firstly, a set of mutated genes specific to lung cancer is selected, and a gene-gene interaction weighted network is constructed using relevant Gene Ontology (GO terms or biological processes). Secondly, a clustering algorithm is applied to detect dense clusters with high weights, resulting in the identification of significant modules associated with lung cancer with high P-values based on cancer-related pathways.

### 2.1 Step I: Gene selection and gene-gene interaction network construction

We construct a lung cancer network with a set of genes with high-frequency mutations associated with lung cancer, with *g*_*i*_ denoting the *i*^*th*^ gene. We extend this set of genes with respect to the biological processes in which each gene *g*_*i*_ participates. Specifically, genes *g*_*i*_ and *g*_*j*_ were connected in the lung cancer network, if and only if, there exists at least one GO term in which both genes participate at the same time. This process leads to the creation of an extended graph of *n* nodes (corresponding to the total number of genes) and *m* edges.

More formally, let *V* = *{g*_1_, …, *g*_*n*_*}* represent the set of lung cancer network genes. For each gene *g*_*i*_, we define 𝒢 𝒪 (*g*_*i*_) as the set of GO terms related to the biological processes in which gene *g*_*i*_ participates. Based on the presence of genes in the GO terms, we define a weighted mutated network *G* =*< V, E, ω >*. The weight of the edge between two connected genes *g*_*i*_ and *g*_*j*_, denoted as *ω*(*g*_*i*_*g*_*j*_), is determined as follows:

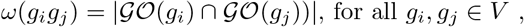

with | *X* | denoting the cardinality or the number of elements in the set *X*. The algorithm to construct the graph for each mutated cancer gene is described in Algorithm 1. In Figure 1, a section of the original graph we obtained in our experimental evaluation is illustrated, specifically showcasing the initial 100 genes along with their corresponding edges.

**Fig. 1.**
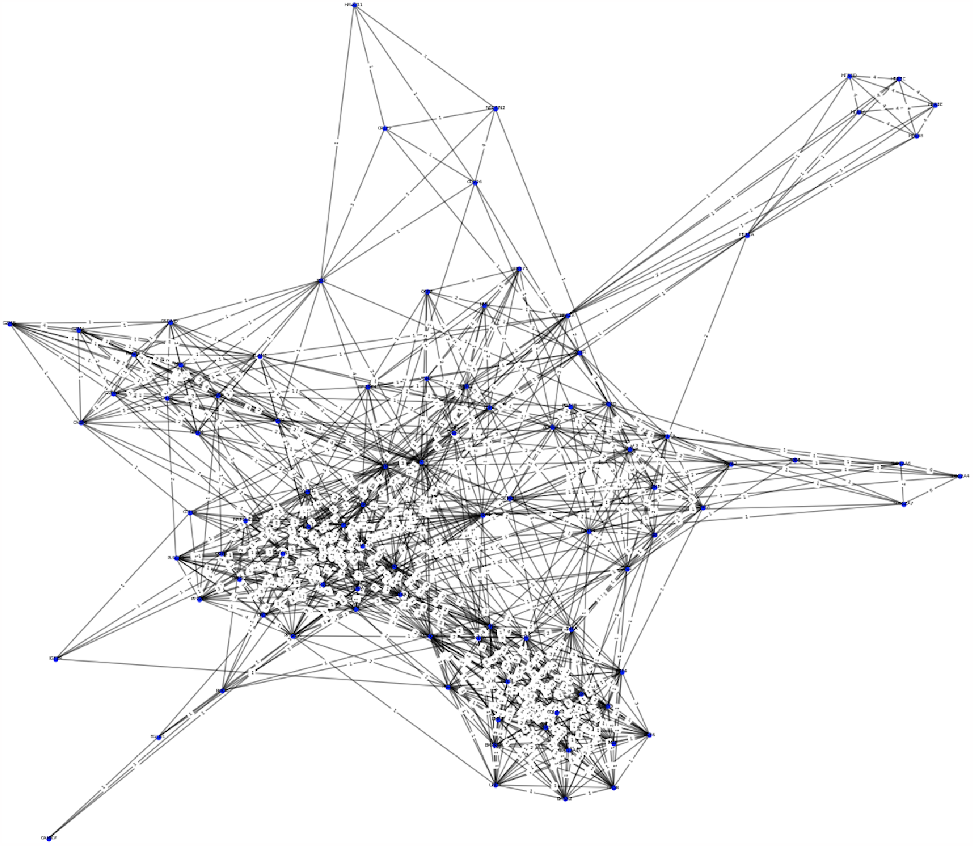
Subgraph of the original gene-gene interaction graph used in our experimental evaluation. In this subgraph we showcase the graph for the first 100 genes.

### 2.2 Step II: Community detection in the gene-gene interaction graph

The second step of our workflow involves the identification of clusters in the gene-gene interaction graph. Without loss of generality, any community detection algorithm can be employed. In this paper, we chose the Leiden algorithm, as it is widely recognized for its effectiveness in community detection. The algorithm optimizes a quality function to identify densely connected groups of nodes [24], and comprises three main phases: (1) local node movement, (2) partition refinement, and (3) network aggregation based on the refined partition. The algorithm initially performs local node movements, followed by refinement of the partition. Finally, the non-refined partition is used to create an initial partition for the aggregate network during the aggregation phase.

Let 𝒢 = (𝒱, ℰ) be a graph with *n* =| 𝒱 | nodes and *m* =| ℰ | edges. Graphs are assumed to be undirected. For simplicity, our mathematical notation assumes graphs to be unweighted but all the mathematical parts are valid for weighted graphs like our constructed graph in the previous section. A partition 𝒫 =*{C*_1_, …, *C*_*r*_*}* consists of *r* = | 𝒫 | communities, where each community *C*_*i*_ *⊆* 𝒱 consists of a set of nodes such that 𝒱 = ⋃_*i*_ *C*_*i*_ and for all *i* / ≠ *j, C*_*i*_ ⋂ *C*_*j*_ ≠ θ.

The Leiden algorithm incorporates a refinement phase to obtain a partition 𝒫_*refined*_, which is a more detailed version of the initial partition 𝒫resulting from the local moving phase. The Leiden algorithm creates an aggregate network based on 𝒫_*refined*_ instead of 𝒫, allowing for the identification of higher-quality partitions. The refinement phase involves merging nodes within communities of 𝒫_*refined*_ while ensuring sufficient connectivity to their communities in 𝒫. This process may split communities in into multiple communities in 𝒫_*refined*_. During the refinement phase, node mergers are not solely based on maximizing the increase in the quality function. Instead, a node can be merged with any community, which leads to an increase in the quality function, with the selection of the community being randomized. The likelihood of selecting a community is influenced by the magnitude of the increase in the quality function. The level of randomness in community selection is controlled by a parameter *θ >* 0, enabling the exploration of a broader partition space. This algorithm (Algorithm 2) offers scalability, enabling the analysis of large graphs that may be computationally demanding for alternative methods. Moreover, the algorithm demonstrates stability by minimizing randomness, ensuring consistent outcomes when applied to the same graph.

#### Algorithm 1

Step I: Constructing the gene-gene interaction graph

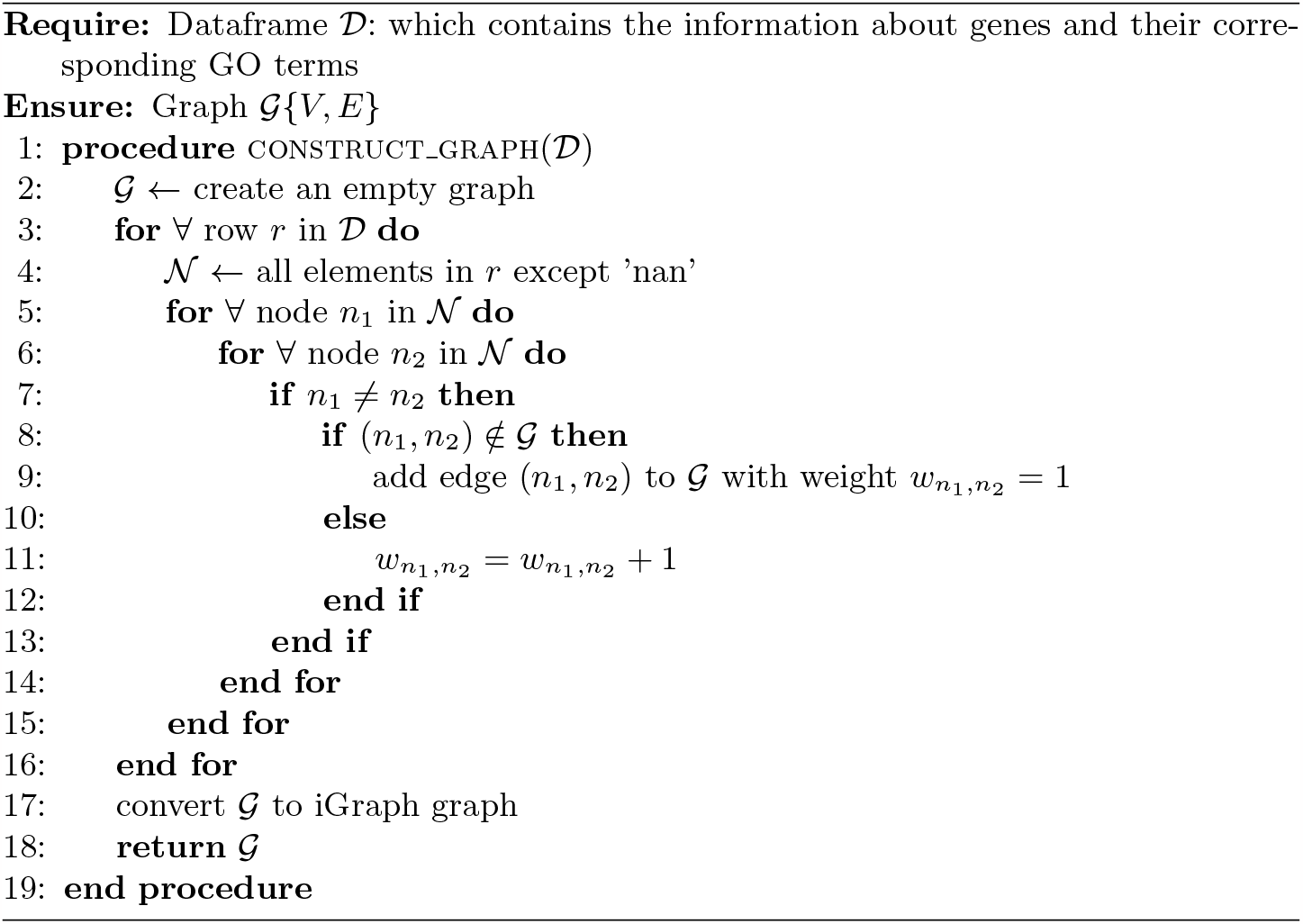

## 3 Experimental results

### 3.1 Datasets

Two different datasets were utilized in this study. The first dataset was used to extract the mutation network, while the second dataset provided the relevant biological process annotations specifically related to lung cancer. All the Materials and implementations are available at our GitHub repository [8].

#### Mutated Genes

The extraction of significantly mutated genes is a crucial step in identifying driver genes and modules associated with lung cancer. To achieve this, researchers have accessed various datasets containing genetic alterations in patients with bronchus and lung cancer. Among these datasets, the TCGA dataset is particularly reliable [9]. We also obtained a dataset comprising mutated genes in lung cancer patients, along with pertinent biological information. From this dataset, we focused on the top 100 genes exhibiting the highest mutation frequency, considering genetic modifications present in multiple cases.

##### Algorithm 2

Step II: Finding communities in the graph

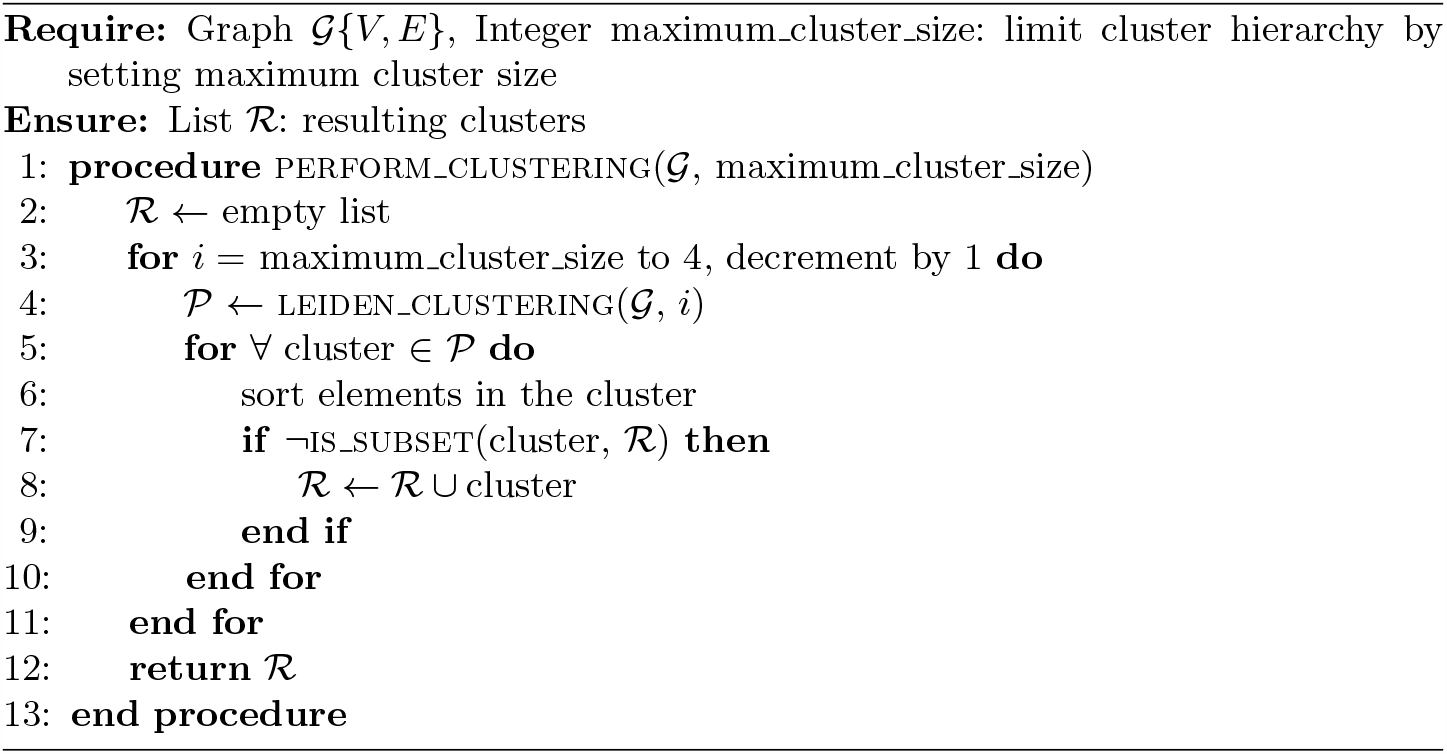

#### Gene Ontology

To identify driver genes and modules in lung cancer, we utilized GO terms from the UniProt database [22]. This extensively curated human-centric database comprises 20,422 entries, providing valuable insights into biological processes and associated proteins. Leveraging the resources within the UniProt database, we conducted a thorough analysis of human proteins, exploring their roles and interactions. We used general GO terms without any filtering, and then we expanded the pool of 100 highly mutated genes for lung cancer with the help of these GO terms. The extended set of genes consists of genes that are functionally related to the mutated lung cancer genes. This approach allowed us to comprehensively explore genes that may not possess genetic alterations but could still contribute to lung cancer.

### 3.2 Setup

The primary objective of this study is to identify modules specifically associated with lung cancer. To achieve this goal, we introduced dense clusters with high weights as potential candidate driver modules related to lung cancer. Our approach involved considering genes within 61 cancer-related biological processes defined by GO terms, resulting in a set of 100 genes with high mutation frequency from TCGA [9]. To identify the candidate set of driver modules related to lung cancer, we applied community detection (using the Leiden algorithm) and subsequently filtered the clusters based on their weights. We generated unique dense clusters of sizes 4, 5, 6, 7, 8, 9, and 10, resulting in a total of 4,694 dense clusters. This specific range is chosen based on the understanding that biological modules typically consist of at least 4 genes, as smaller groupings may lack the necessary complexity and functionality associated with modular systems in biology [23, 25, 26]. Conversely, clusters with more than 10 genes are considered excessively broad and tend to provide less informative insights. Their larger size compromises their specificity, making it challenging to discern the distinct functional roles of individual genes within the broader biological context [25]. Thus, by focusing on clusters within the 4–10 member range, we aim to strike a balance between complexity and specificity, facilitating a more nuanced understanding of gene interactions and their functional significance. Furthermore, our approach focuses on the identification of clusters that are not part of any other cluster. The objective is to uncover distinct and exclusive groups of genes that exhibit unique characteristics or functionalities. By doing so, we aim to identify exceptional clusters that possess specific properties or perform specialized functions, separate from the influence of larger clusters. This approach allows us to uncover novel insights into the intricate dynamics of the genetic network by uncovering hidden patterns and novel insights into the underlying biology of lung cancer modules. Next, from the obtained dense clusters, we selected the top 5% of clusters with the highest average weight, which are considered as the candidate driver module set. The candidate driver modules are dense clusters and good clusters in the sense of clustering quality. We showed them in 7 tables as supplementary materials in our GitHub repository [8]. In this study, we are looking for biologically meaningful modules related to lung cancer; therefore, we defined the P-value score to measure the association of these modules with cancer-related pathways, and then we selected 110 modules from this candidate set based on the P-value of the cancer-related pathways associated with each module. These modules represent subsets of genes that demonstrate significant involvement in cancer pathways and we presented these filtered modules in Tables 1–7, respectively.

**Table 1.** Statistically significant lung cancer driver modules of size 4.

| Genes within the module |  |  |  | P-values corresponding to the signaling pathways implicated in cancer development |  |  |  |  |  |  |  |  |  | # of P-values below the 0.05 threshold |
| --- | --- | --- | --- | --- | --- | --- | --- | --- | --- | --- | --- | --- | --- | --- |
|  |  |  |  | ERBB | Estrogen | FoxO | MAPK | mTOR | p53 | PI3K | Ras | VEGF | Wnt |  |
| CDKN2A | CDKN2B | CDKN1A | HRAS | 0.000 | 0.034 | 0.000 | 0.074 | 0.041 | 0.000 | 0.003 | 0.058 | 0.015 | 0.956 | 7 |
| EGFR | ID1 | EGF | ERBB2 | 0.000 | 0.034 | 0.000 | 0.000 | 0.959 | 0.981 | 0.000 | 0.001 | 0.984 | 0.956 | 6 |
| GSK3B | GSK3A | CSNK1A1 | CSNK2A3 | 0.022 | 0.965 | 0.965 | 0.923 | 0.041 | 0.981 | 0.086 | 0.941 | 0.984 | 0.000 | 3 |
| FGFR2 | WNT2 | FGF9 | TMIE | 0.977 | 0.965 | 0.965 | 0.002 | 0.041 | 0.981 | 0.003 | 0.001 | 0.984 | 0.044 | 5 |
| NODAL | SP3 | CITED2 | JUN | 0.022 | 0.034 | 0.965 | 0.074 | 0.959 | 0.981 | 0.911 | 0.941 | 0.984 | 0.044 | 3 |
| FST | FGF10 | FGF7 | NKX2-1 | 0.977 | 0.965 | 0.965 | 0.002 | 0.959 | 0.981 | 0.003 | 0.001 | 0.984 | 0.956 | 3 |
| SMPD3 | PDGFB | PDGFA | NR4A3 | 0.977 | 0.965 | 0.965 | 0.002 | 0.959 | 0.981 | 0.003 | 0.001 | 0.984 | 0.956 | 3 |
| CSF1R | CSF1 | IL34 | DOK7 | 0.977 | 0.965 | 0.965 | 0.002 | 0.959 | 0.981 | 0.003 | 0.001 | 0.984 | 0.956 | 3 |
| TP53 | TIGAR | MDM2 | RPL26 | 0.977 | 0.965 | 0.034 | 0.074 | 0.959 | 0.000 | 0.003 | 0.941 | 0.984 | 0.044 | 4 |
| MAPK14 | MAP3K20 | MAPK11 | DYM | 0.977 | 0.965 | 0.000 | 0.000 | 0.959 | 0.981 | 0.911 | 0.941 | 0.000 | 0.956 | 3 |
| BCL2 | IL7 | IL9 | RAG2 | 0.977 | 0.034 | 0.034 | 0.923 | 0.959 | 0.019 | 0.003 | 0.941 | 0.984 | 0.956 | 4 |
| PIK3CG | GPR183 | PIK3CD | TREM1 | 0.022 | 0.034 | 0.034 | 0.923 | 0.041 | 0.981 | 0.003 | 0.058 | 0.015 | 0.956 | 6 |

### 3.3 Evaluation of modules related to lung cancer

Extensive research has unraveled the intricate signaling pathways involved in cancer development. Disruption of these pathways is critical to driving cancer progression. Building upon Ahmad et al. [27], we have compiled a comprehensive list of cancer-related signaling pathways, which provides valuable insights into the molecular mechanisms underlying cancer progression. These pathways govern crucial cellular responses like survival, proliferation, migration, differentiation, and apoptosis, forming the foundation of our investigation into lung cancer-specific modules.

#### ERBB signaling pathway

The ERBB family plays a crucial role in cancer-related signaling pathways, including proliferation, survival, angiogenesis, and metastasis. These receptors are frequently amplified, mutated, or overexpressed in cancer cases, making them attractive targets for cancer treatment. Dysregulation and mutations within the ERBB family have been found to play a significant role in evading anti-tumor immune responses, further emphasizing their importance in cancer progression [27].

#### Estrogen receptor signaling pathway

Estrogen’s involvement in lung cancer has been studied [29]. Recent research has explored its impact on lung cancer development, signaling pathways, and the connection between estrogen receptor (ER) and epidermal growth factor receptor (EGFR). Clinical trials combining ER and EGFR antagonists have been discussed. The detrimental effects of tobacco smoking on estrogen and the role of environmental endocrine-disrupting chemicals targeting ER in lung carcinogenesis are also significant considerations.

#### FOXO signaling pathway

FOXOs are crucial players in cell fate decisions and tumor suppression across various cancer types. The PI3K/AKT pathway is a key player in the interaction with FOXOs and is implicated in several types of cancer. Additionally, other pathways such as Ras-MEK-ERK, IKK are associated with FOXOs in tumorigenesis.

#### MAPK signaling pathway

The MAPK pathway is a tightly controlled part that plays a vital role in diverse cellular processes, including migration, cell proliferation, differentiation, and survival [30]. Dysregulation of this pathway has been implicated in various types of cancer. Additionally, parallel or redundant pathways, such as the PI3K/Akt pathway, are frequently disrupted in cancer.

#### mTOR signaling pathway

The mTOR pathway regulates crucial cellular processes like cell survival, metabolism, growth, and protein synthesis. Dysregulation of mTOR signaling frequently occurs in various malignancies, including lung cancer [31]. Hyperactivation of the mTOR pathway promotes cell proliferation and metabolism, contributing to tumor initiation and progression. Furthermore, mTOR signaling negatively regulates autophagy through multiple mechanisms.

#### P53 signaling pathway

p53 gene mutations are commonly found in human neoplasms, including lung cancer, and are linked to aggressive tumor characteristics and lower overall survival rates. Acting as a tumor suppressor, p53 plays a vital role in maintaining cancer control and inhibiting abnormal cell growth [32]. Post-translational modifications, like ubiquitination, are key in regulating the activity and stability of wild-type p53.

#### PI3K/Akt signaling pathway

The EGFR-mediated signaling pathway includes the PI3K/AKT pathway, which is frequently dysregulated in various human malignancies [28]. Extensive research has shown that PI3Ks are crucial in regulating essential cellular processes involved in cancer development, including metabolism, cell survival, proliferation, differentiation, and motility [28]. The PI3K/AKT pathway is governed by multiple oncogenes and growth factor receptors, such as KIT, MET, EGFR, and ERBB, playing a significant role in intracellular physiological processes [27].

#### Ras/Raf signaling pathway

Targeting the Ras/Raf signaling pathway and its upstream activators has gained attention in cancer research. Abnormal activation of this pathway, particularly through EGFR activation, is highly prevalent [27]. Ras proteins, central to this pathway, are frequently implicated in oncogenesis, with their mutations associated with cancer development [28].

#### VEGF signaling pathway

VEGF signaling in lung cancer extends beyond angiogenesis, impacting various aspects of tumor progression. This pathway enables carcinoma cells to evade apoptosis and advance toward invasive and metastatic states [33]. Cells with active VEGF signaling gain a survival advantage and promote lung cancer dissemination. These findings emphasize the potential of targeting VEGF and its receptors to directly address tumor cells.

#### Wnt signaling pathway

The Wnt signaling pathway holds a crucial role in the regulation of embryonic organ development and cancer progression. Emerging studies have shown its importance in various aspects, including therapeutic resistance, maintenance of stemness, regulation of the immune microenvironment, and shaping of the cancer phenotype [34].

For evaluation, we used the Gene Set Enrichment Analysis (GSEA) formula to calculate the P-value of candidate modules with respect to the mentioned cancer-related pathways. Suppose that *P* = {*p*_1_, …, *p*_*M*_} is a collection of the above-mentioned pathways related to cancer and *H* is a candidate module (cluster) for lung cancer. The GSEA equation is employed to determine the P-value of cluster *H* in relation to pathway *P*_*i*_, described as follows:

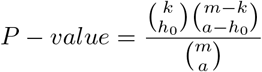

In this formula, *m* represents the total genes present in the lung cancer network and *a* is the number of genes in the cluster. Furthermore, *k* shows the number of genes in pathway *p*_*i*_, and *h*_0_ indicates the number of genes in the cluster found within the assumed pathway. A cluster was identified as a lung cancer driver module if it exhibited a significant P-value, high density, and an elevated average score. A threshold of 0.05 was set for the P-value, and a module was identified as a cancer driver if it had a P-value below this threshold for at least three distinct cancer-related pathways.

### 3.4 Results

The results are presented in this subsection in 7 tables according to the size of the modules and their corresponding P-value in each of the cancer-related signaling pathways. In all tables, the first column indicates the list of gene names, columns 2–11 display the corresponding P-values for each selected signaling pathway and the last column indicates the number of signaling pathways in which the module’s P-value falls below the 0.05 threshold.

Table 1 shows the significant modules with 4 genes and emphasizes the significance of proposed clusters in terms of their biological relevance. The presence of multiple modules showing statistical significance concerning the predefined pathways supports this observation. We successfully identified 12 distinct clusters, each consisting of 4 genes, and a considerable proportion of these clusters exhibited statistical significance across three or more pathways, as indicated in the last column. Similarly, Table 2 presents the statistically significant modules consisting of 5 genes. Unlike Table 1, Table 2 exhibits a distinct characteristic, as it contains several modules that show statistical significance for eight different pathways. This represents the highest number of pathways identified for any cluster size, although this feature is not exclusive to Table 2. Table 3 provides an overview of modules comprising 6 genes that demonstrate statistical significance. This table contains several modules that show statistical significance for seven different pathways. Table 4 shows lung cancer driver modules consisting of 7 genes. This table contains several modules that show statistical significance for more than five different pathways. Table 5 displays the modules of size 8. Notably, the gene PDGFB appears most frequently, appearing in a total of 7 clusters. It is followed by CDKN1A, INS, INSR, IGF1R, and AKT1, which each appear in 6 clusters. Among the various cluster sizes explored, as evident in Table 5, which contains the highest number of modules (a total of 22). Table 5 contains several modules that show statistical significance for eight different pathways. Table 6 presents the lung cancer driver modules, consisting of 9 genes. This table contains several modules that show statistical significance for seven different pathways. Table 7 illustrates the largest modules, consisting of 10 genes. This table contains several modules that show statistical significance for more than five different pathways. These modules provide a foundation for further investigations, such as exploring potential combinations between modules and examining the implications of their statistical assessment.

**Table 2.**
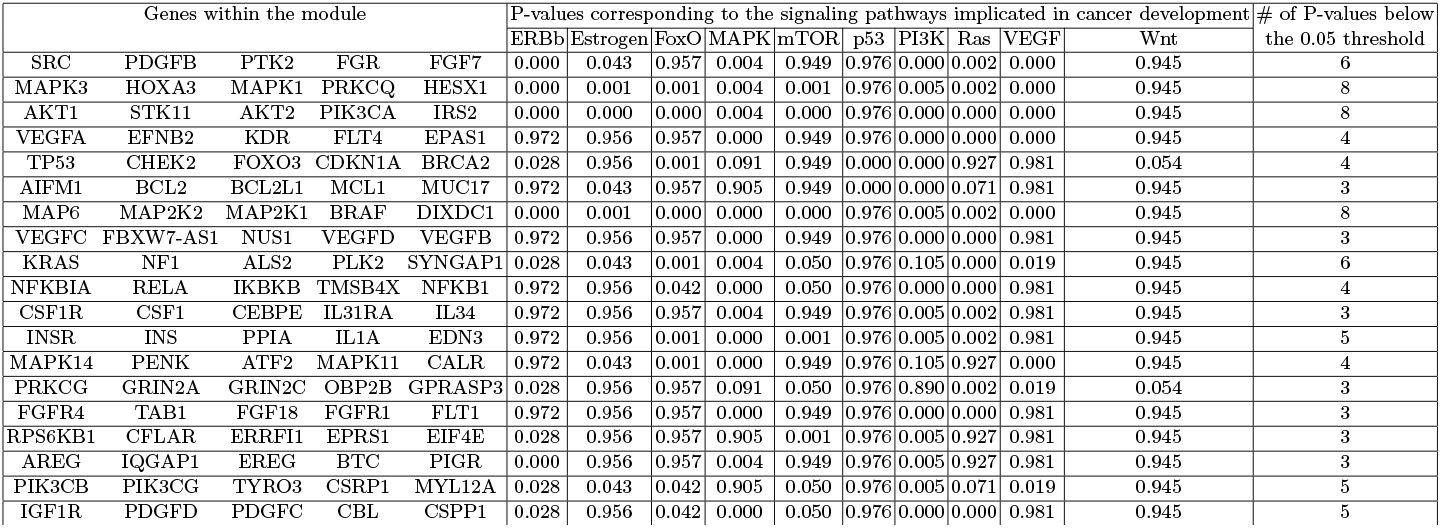
Statistically significant lung cancer driver modules of size 5.

**Table 3.**
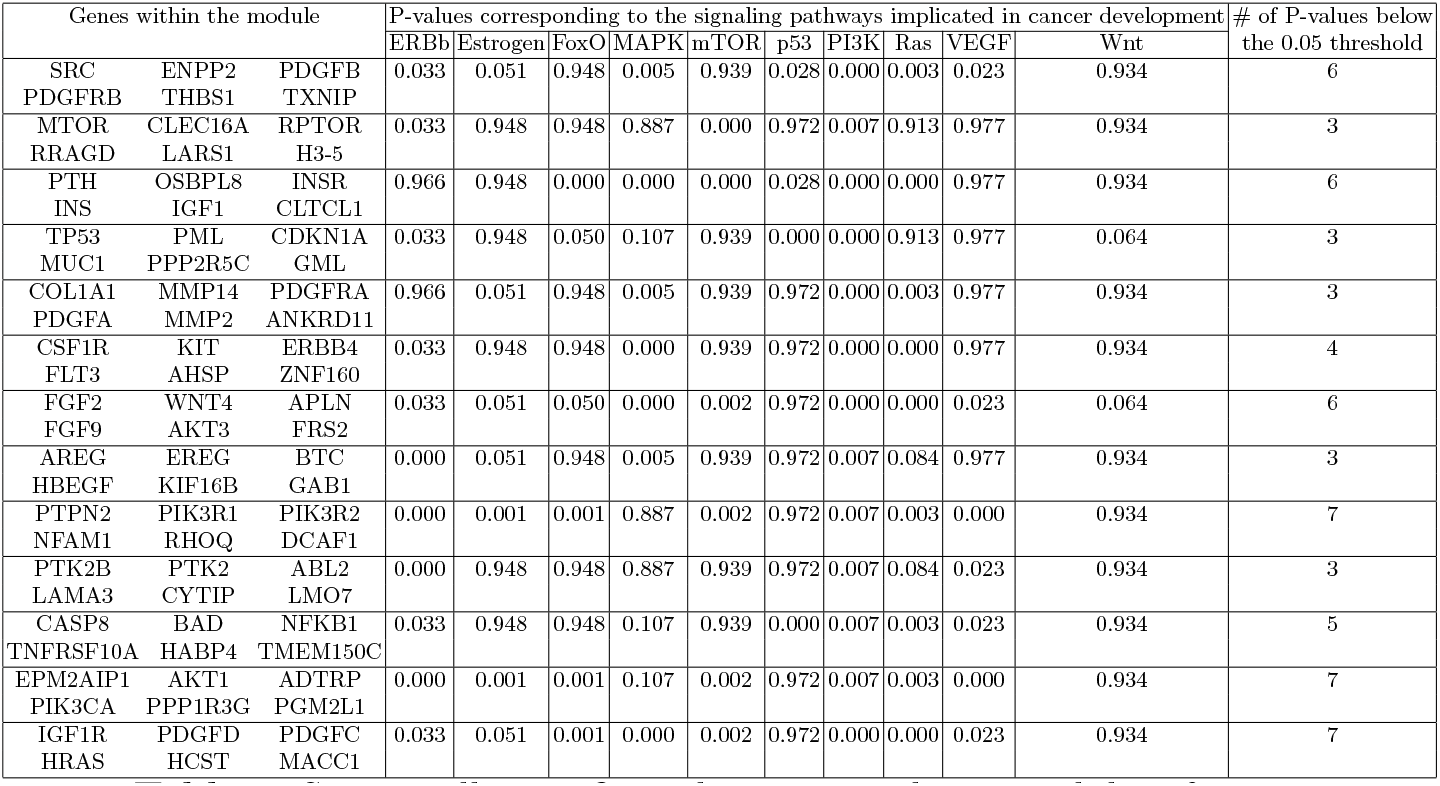
Statistically significant lung cancer driver modules of size 6.

**Table 4.**
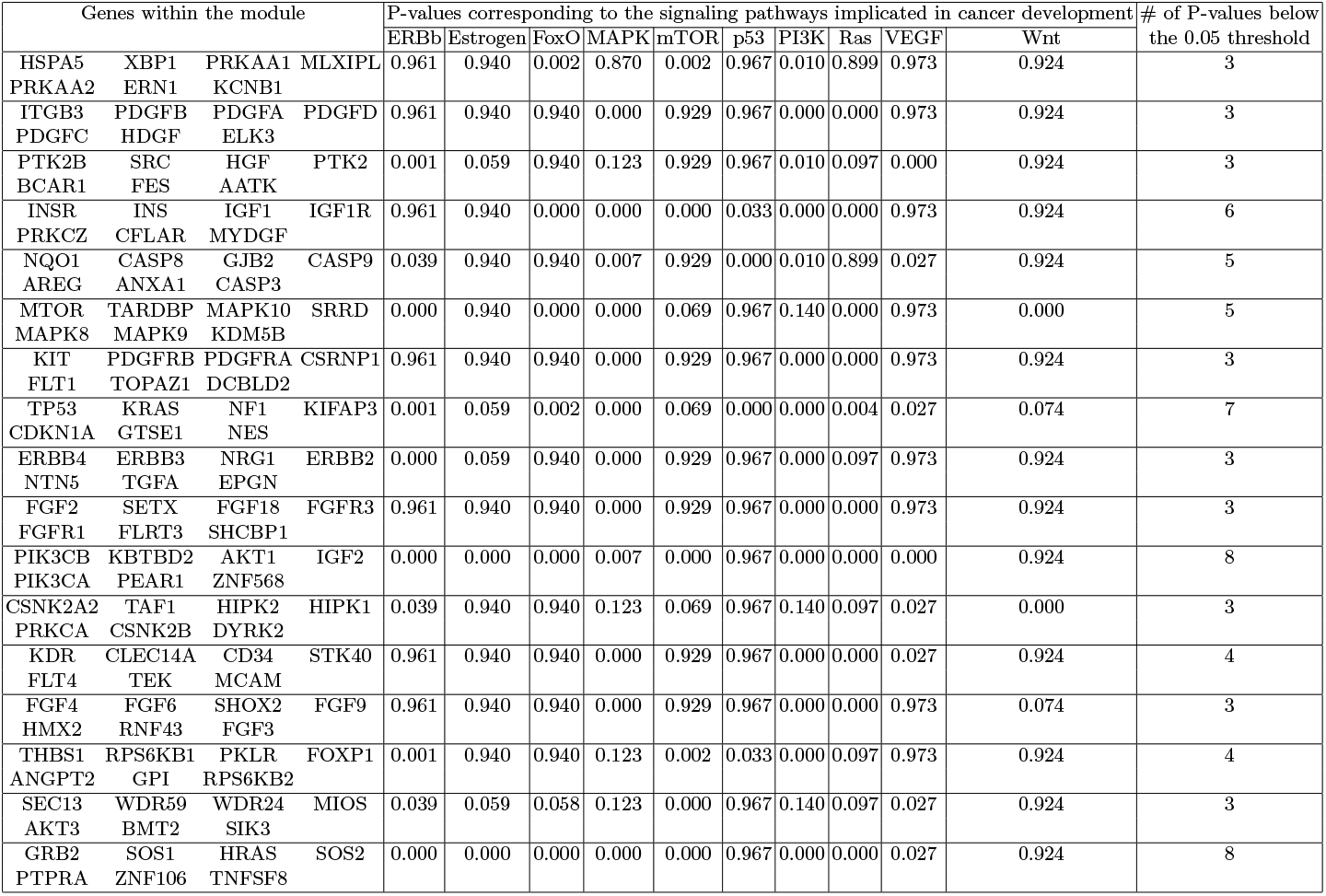
Statistically significant lung cancer driver modules of size 7.

**Table 5.**
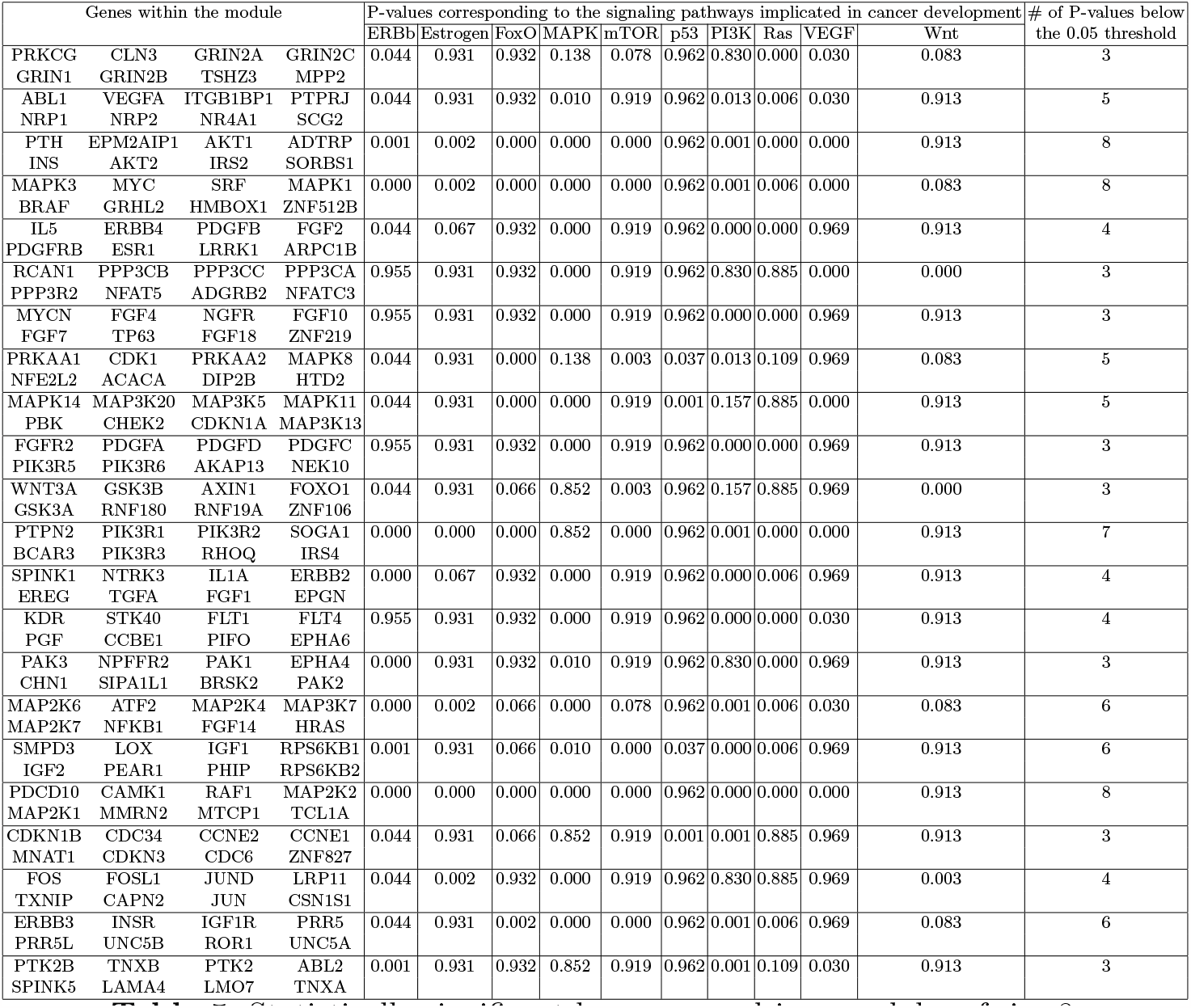
Statistically significant lung cancer driver modules of size 8.

**Table 6.**
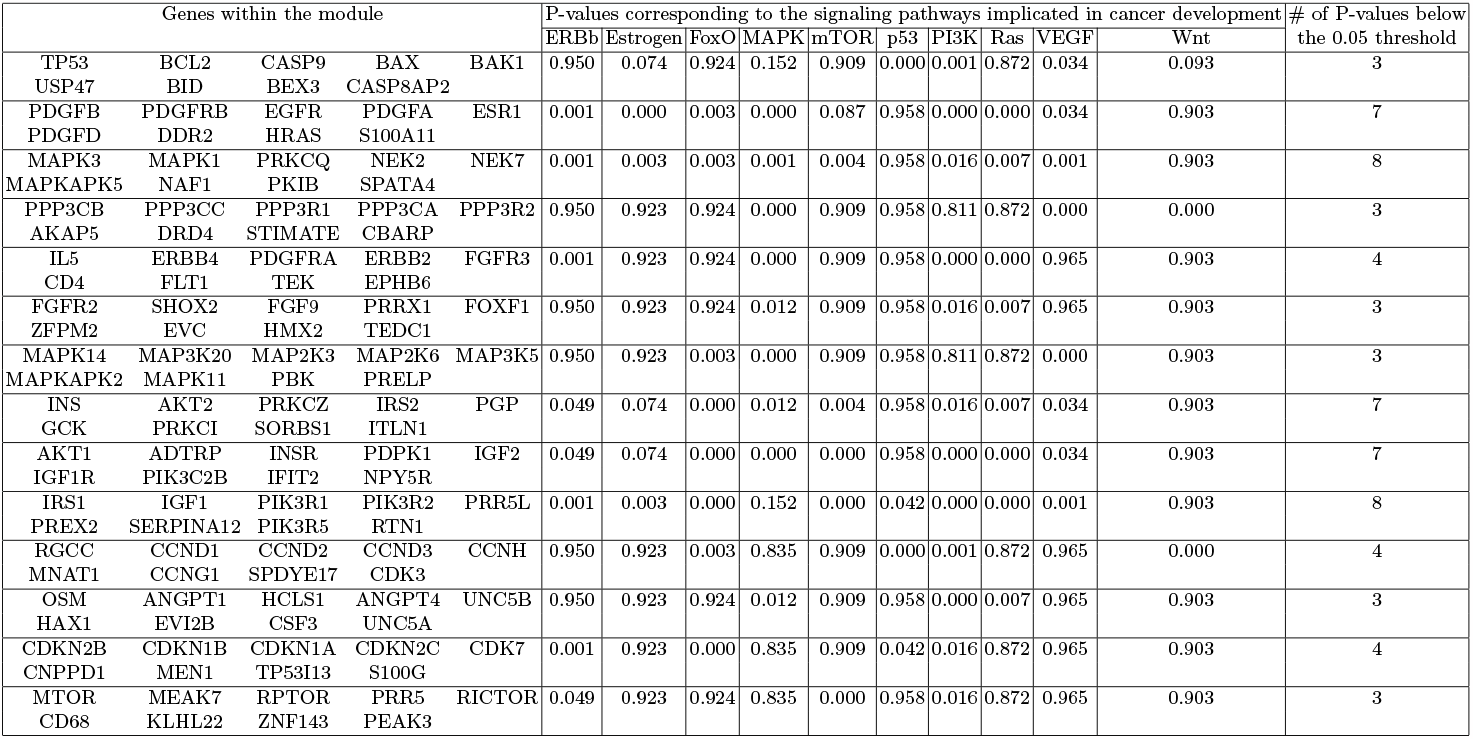
Statistically significant cancer driver modules of size 9.

**Table 7.**
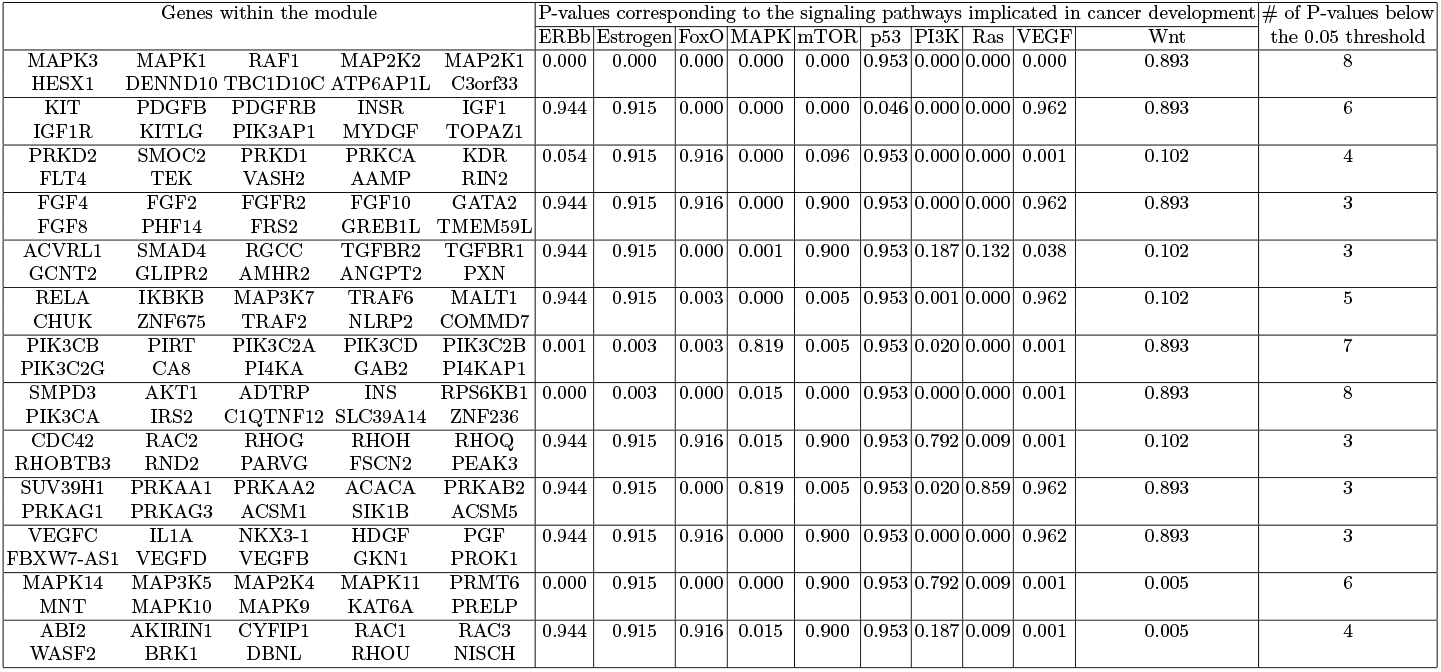
Statistically significant cancer driver modules of size 10.

In total, this work identified 110 cancer-driver modules that are both statistically and biologically significant. Among these, 46 modules are associated with 3 different pathways, 18 modules with 4 pathways, 11 modules with 5 pathways, 13 modules with 6 pathways, 10 modules with 7 pathways, and 12 modules with 8 pathways. These findings highlight the diversity and complexity of the molecular mechanisms involved in cancer development and progression. It is worth noting that out of the 10 pathways analyzed, the PI3K pathway demonstrated the highest statistical significance with P-values below the 0.05 threshold for a total of 89 or 80.9% of all modules. On the other hand, the Wnt pathway showed the least statistical significance, appearing in only 13 modules, which corresponds to 11.8% of all modules generated by this clustering method. Figure 2 provides a comprehensive visualization, presenting the P-values associated with each module within various pathways. The heatmap offers valuable insights into the statistical significance and relative strengths of these modules.

**Fig. 2.**
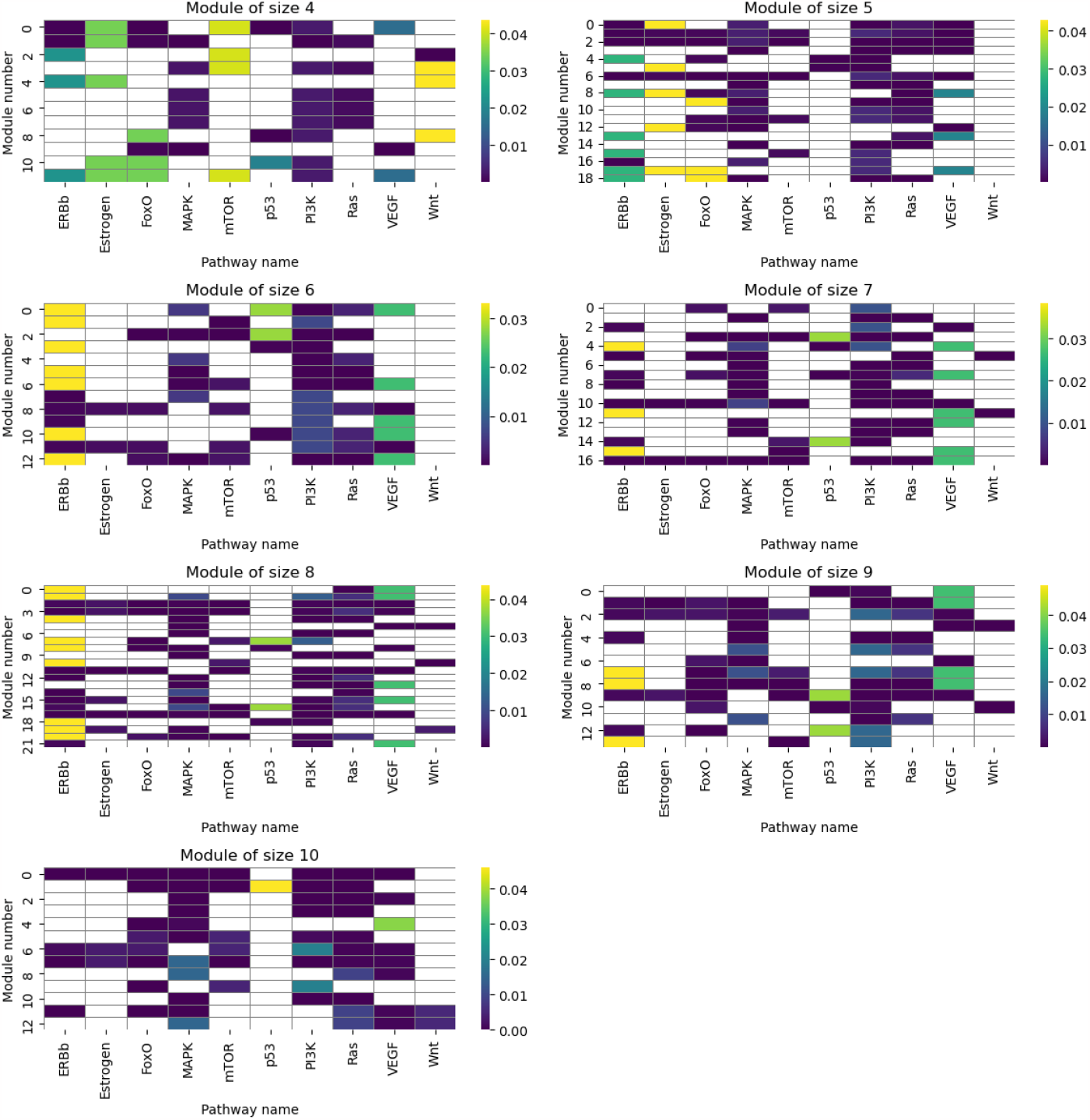
The heatmap, shows the intricate relationships between the modules and their corresponding pathway P-values

In summary, the analysis of the 110 distinct modules identified has revealed their statistical significance across a minimum of three selected signaling pathways. Remarkably, a majority of these modules surpass this minimum requirement, demonstrating statistical relevance in up to eight pathways. The interconnected nature of each module suggests their involvement in multiple biological processes, with no module being a subset of another. These findings are highly satisfactory, providing substantial insights into the field of lung cancer research and highlighting the significance of these modules in understanding the disease.

## 4 Conclusions

Lung cancer, a leading cause of cancer-related mortality globally, poses a significant challenge in terms of treatment due to its genetic alterations and heterogeneity. In this study, our focus was on identifying driver modules associated with lung cancer. By utilizing a collection of mutated genes found in lung cancer patients, we identified the key genes responsible for driving lung cancer. To construct a biological network that represents the mutated gene set, we assigned network nodes to the mutated genes, and the network edge between the two genes shows the number of GO terms that both genes participated in at the same time. Next, we introduced a clustering method as a machine-learning approach. This algorithm facilitated the identification of high-density sub-networks with significant weight within the lung cancer network. We discovered the top 5% of clusters with the highest average weight and calculated the P-value for each of these obtained modules based on well-established cancer pathways. Consequently, we presented 110 significant modules in Tables 1-7 showcasing their extensive coverage within cancer pathways. The identification of driver genes and modules in lung cancer offers a comprehensive understanding of the causes and progression of this disease.

